# Environmental, structural and taxonomic diversity factors drive aboveground carbon stocks in a semi-deciduous tropical rainforest strata in Cameroon

**DOI:** 10.1101/2021.03.12.435155

**Authors:** Jules Christian Zekeng, Masha T. van der Sande, Jean Louis Fobane, Reuben Sebego, Wanda N. Mphinyane, Paul André Ebanga, Marguerite Marie Abada Mbolo

**Affiliations:** Department of Plant Biology, Faculty of Science, University of Yaounde I, PO Box: 812 Yaounde, Cameroon; Department of Environmental Science, Faculty of Science, University of Botswana, Private Bag UB 0704 Gaborone, Botswana; Department of Biological Sciences, Florida Institute of Technology, Melbourne, FL, USA; Institute for Biodiversity & Ecosystem Dynamics, University of Amsterdam, Amsterdam, The Netherlands; Forest Ecology and Forest Management Group, Wageningen University and Research, Wageningen, The Netherlands; Department of Biological Sciences, Higher Teachers’ Training College, University of Yaounde I, PO Box 47, Yaounde, Cameroon

**Keywords:** biodiversity, logging disturbance, ecosystem functioning, forest structure, soil fertility/texture, topography

## Abstract

Tropical forests play an important role in biodiversity conservation and ecosystem functioning. Few studies have teased apart the independent effects of biotic and abiotic factors on aboveground carbon across tree size groups and the whole tree community level in a tropical semi-deciduous rainforest. This study aims to analyze the relative and independent effects of abiotic (topography, soil fertility/texture and disturbance) and biotic factors on carbon stocks across tree size groups, as well as at the whole tree community in semi-deciduous plots, including logged plots.

We used data from 30 1-ha plots and 22,064 trees distributed in a semi-deciduous tropical rainforest of Cameroon. For each plot, we quantified disturbance, topography and eleven soil conditions. Besides, we quantified three taxonomic richness and diversity Gini index. We used structural equation models to test the hypothesis that all drivers have independent, positive effects on aboveground carbon stocks across tree-sized groups and the whole tree community.

The logging disturbance decreased aboveground carbon stocks for the whole tree community and trees size groups across our site except for large trees. Topographical factors (i.e., slope and elevation) increased aboveground carbon (AGC) of the whole tree community and the large trees. In contrast, soil nitrogen (Nsoil) and clay proportion increased AGC of the small stems and understorey trees groups. This study found that biotics factors have some indirect and direct effects on AGC. Taxonomic diversity through niche complementarity had a positive relationship with aboveground carbon stocks across the trees size groups and the whole tree community, while diversity Gini index had strongest relationships with aboveground carbon stocks.

The significant effects of structural diversity on aboveground carbon stocks of the whole tree community, large trees, and understory trees highlight the importance of maintaining a layered structure and tall trees in the forest.

## 1. Introduction

Tropical forests are at the center of any existing debate on climate change and sustainable forest management because of their dual roles in climate change mitigation and biodiversity conservation (Arasa-Gisbert *et al*., 2018; Bele *et al*., 2015; Bodegom *et al*., 2009; Poorter *et al*., 2016). It has been shown that biodiversity can enhance forest productivity and carbon storage in Neotropical forests (Poorter *et al*., 2017), suggesting that climate change mitigation and biodiversity conservation go hand in hand. Here, we assess how biodiversity and forest structure (as biotic factors) and abiotic factors interactively determine forest carbon storage in semi-deciduous tropical rainforests in Cameroon.

Earlier studies have assessed the simultaneous effects of abiotic (e.g. soil fertility) and biotic (e.g., biodiversity and stand structure) factors on forest functioning based on trees with diameter ≥ 10 cm (Fotis *et al*., 2018; Poorter *et al*., 2017; van der Sande *et al*., 2018; van der Sande *et al*., 2017), but few have done that for smaller trees and understorey vegetation. Moreover, most of these studies have been realized in Neotropical forests (Chisholm *et al*., 2013; Poorter *et al*., 2017; van der Sande, 2016; van der Sande *et al*., 2018) and subtropical Asiatic forest (Ali *et al*., 2019; Ali & Yan, 2017). Understory vegetation and small trees grow in lower light levels and may consist of different species composition. For that reason, processes determining carbon stocks of large trees may not be the same as processes determining carbon stocks of small trees and understory vegetation. Here, we evaluate how abiotic and biotic factors, directly and indirectly, affect aboveground carbon stocks of different tree size groups and the whole tree community in a Cameroonian semi-deciduous forest.

Taxonomic diversity and richness can affect carbon stocks (Fig. 1) through a variety of mechanisms: (1) niche complementary or facilitation among species is thought to be a key mechanism by which biodiversity affects the rates of resource use that govern the efficiency and productivity of ecosystems (Tilman *et al*., 2001); (2) the selection effect hypothesis suggesting that diversity effects are caused by a greater chance of one or a few dominant, high biomass species being present in the community (Loreau & Hector, 2001); (3) insurance effect, where more diverse communities have been shown to have higher and more temporally stable ecosystem functioning than less diverse ones, suggesting they should also have a consistently higher level of functioning over time (Allan *et al*., 2011).

**Fig 1.**
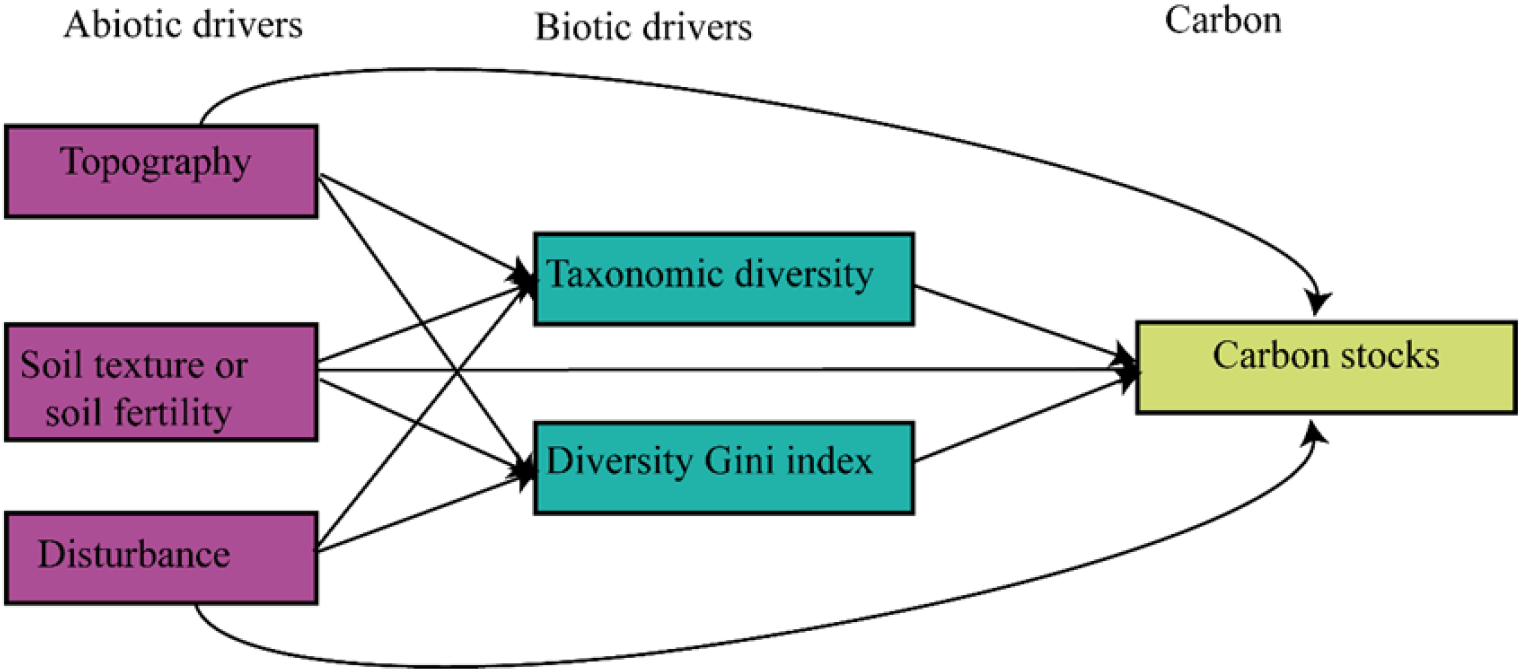
Conceptual framework linking abiotic drivers (topography, soil texture/fertility and disturbance) and biotic drivers (taxonomic diversity and structural attributes) to carbon stocks. Disturbance is included as abiotic drivers because it affects forest density and light availability.

Besides taxonomic attributes, forest structure, such as stem diameter, tree density and structural diversity, can determine biomass storage (Ali *et al*., 2019; Fotis *et al*., 2018; Poorter *et al*., 2015). Forest structure influences plant light capture and usage (Laurans *et al*., 2014), thereby shaping aboveground biomass between overstorey and understorey strata.

Local abiotic factors (e.g., climate, soils, topography) can both directly and indirectly (via biotic factors) affect aboveground biomass (Poorter *et al*., 2017; van der Sande *et al*., 2018). For example, steep slopes decrease, and increasing elevation decreases biomass stocks (Gonmadje *et al*., 2017). Earlier studies in the Congo basin have shown that altitude (Gonmadje *et al*., 2017), soil nutrients (Doetterl *et al*., 2015; Fayolle *et al*., 2012; Lewis *et al*., 2013), and biotic factors (Bastin *et al*., 2018; Fayolle *et al*., 2016; Zekeng *et al*., 2020) are essential drivers of aboveground biomass. The heterogeneity of soil types and topography at small spatial scales influence soil conditions. Indeed, soil fertility has been shown to positively affect biomass stocks in tropical forests such as on the old and leached nutrient-poor soils of the Guiana shield (van der Sande *et al*., 2018) and of the Doume Communal Forest in Cameroon (Zekeng *et al*., 2020).

Disturbances may modify the vegetation by removing biomass and opening up the forest canopy, leading to the increased availability of light and other resources (Toledo *et al*., 2012), thereby promoting the growth of the remaining trees and hence in the long term, overcompensate for the loss in growth from removed trees (van der Sande *et al*., 2018). Depending on the frequency, intensity, and type, disturbances bring modification in habitat heterogeneity, shifts in competitive balances among species, and the creation of rare habitats, thereby improving species diversity (Denslow, 1995). Where disturbance is occasional, older and larger trees should dominate, reducing the growth rates and survival probability of small trees. Therefore, such low disturbance would lead to more biomass allocated to fewer stems (Holm *et al*., 2014; Lewis *et al*., 2013).

This study aims to analyze the relative independent effects of abiotic and biotic factors on aboveground carbon stocks across tree size groups: small stems (< 5 cm DBH), understorey trees (10 >cm DBH< 5), large trees (> 10 cm DBH), and the whole tree community. We address two questions. Question 1: how do taxonomic diversity (i.e., rarefied species richness, Shannon-Weaver index, species richness) and diversity Gini index influence carbon stocks of each tree size group and the whole tree community? We hypothesize that high species diversity positively affects carbon stocks (through niche complementarity, the selection effect, or the insurance effect) and that diversity Gini index increases carbon stocks. Question 2: how do abiotic conditions influence carbon stocks directly and indirectly via taxonomic diversity and diversity Gini index? We hypothesize that carbon stocks increase with soil fertility or texture and that carbon stocks at trees size groups or the whole tree community most strongly and positively affected by topographic factors. We also hypothesize that longtime disturbance will decrease trees’ carbon stocks with DBH ≥ 5 cm while increasing carbon stocks of small stems.

## 2. Material and Methods

### 2.1. Research site

The research was carried out in the moist, semi-deciduous forest of Doume Communal, eastern Cameroon (4° 31’0” S, 13° 47’5” W). This site receives between 1300 and 1800 mm rainfall per year, with a dry season from November until March. The forest is located on ferralitic red, loose and permeable soils. These soils are poor in nutrients, aciditic and fragile. In the shallows, the soils are hydromorphic to gley. The relief of the forest can be described as slightly uneven. It presents a succession of low hills with generally gentle slopes interspersed with small well-marked streams, or swampy depressions (several hundred meters) without a distinct watercourse (Anonymous, 2015). The altitude varies from 605 to 760 m, with some particularly marked summits, culminating at less than 700 m of altitude.

### 2.2. Sampling plots and sample design

Previous studies using remote sensing and geographical information systems defined the type of land use and land cover in the study site (Zekeng *et al*., 2019). As a result, it allowed us to choose to work only on the terra-firme forest while avoiding rivers and swampy vegetation types. Thirty 1-ha (100 x 100 m) plots were set up in the Doume Communal forest across four villages. The 1-ha plots were subdivided into 25 20 x 20 m subplots. In the whole 1-ha plot, trees ≥ 10 cm diameters at breast height (DBH) (hereafter ‘large trees’), and in thirteen subplots, trees with DBH between 9.9 and 5.0 cm (hereafter ‘understorey trees’) were identified and measured. In the subplots situated in the four corners and the center of each 1-ha plot, a quadrat of 5 m x 5 m was installed to inventory trees between 1.0-4.9 cm diameter at 30 cm aboveground (hereafter ‘small stems’). Hence, in total per 1-ha plot, we sampled all trees ≥ 10 cm DBH, trees between 5 and 10 cm DBH in 5200 m^2^, and trees between 1 and 5 cm diameter in 125 m^2^.

### 2.3. Carbon stock estimation

We calculated carbon stocks of the whole tree community (all trees ≥ 1 cm diameter) and the different tree size groups (i.e. small stems, understorey trees and large trees). Their biomass was converted to carbon using conversion factors according to the recommendation of IPCC (2006): a conversion factor of 0.47 (Thomas & Martin, 2012) widely used in the literature review was used.

Large and understorey trees were measured at 1.3 m breast height or, if applicable, 50 cm above the top of the buttresses or 2 cm above the deformity (Condit, 1998) while small stems were measured at 0.30 m aboveground level. We calculated aboveground biomass (AGB) for large and understorey trees using Eq. (1) of Chave *et al*. (2014) but see Réjou-Méchain *et al*. (2017). The aboveground biomass of small stems was computed using Eq. (2) developed by Djomo & Chimi (2017).

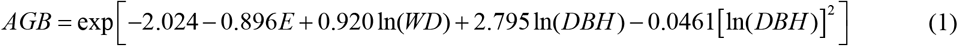

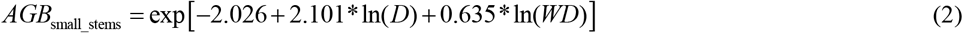

Where E is a measure of environmental stress of the site, which depends on temperature seasonality and water deficit and is extracted from http://chave.ups-tlse.fr/pantropical_allometry/readlayers.r with the retrieve_raster function in R. DBH is the diameter at breast height (cm), and WD is the wood density (g cm^-3^). WD was based on local wood density if available, and otherwise on wood density obtained from the Global Wood Density Database (Chave *et al*., 2009; Zanne *et al*., 2009). Species-level WD was used for 61.5% plots species while genus or family-average per plot were used for 31.9% of species. For the few cases (forty-six species) without genus- or family-level WD (5.6%), we used WD averaged per plot.

### 2.4. Taxonomic diversity

For the whole tree community and each tree’s size groups per 1-ha plot, three species diversity measures were computed (Appendix S1): species richness (number of species per plot), rarefied species richness, and Shannon-Weaver index (Shannon & Weaver, 1949). Rarefied species richness is the number of species observed when a fixed number of trees are randomly drawn from a plot, there-fore removing the confounding influence of tree density on species richness (Poorter et al. 2017). We calculated rarefied species richness here as the number of species at a random draw of 469 stems for the whole tree community, 458 stems for large trees, 62 stems for understorey trees and ten for small stems, as these numbers of individuals were found in all 1-ha plots according to the sampling design. The Shannon-Weaver index requires species abundance and was calculated as follows: 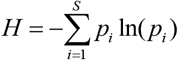, Where P_i_ is the proportion of individuals belonging to the i^th^ species found in a sample. The calculations were done using the R package vegan (Oksanen *et al*., 2018).

### 2.5. Structural diversity

This study aimed to determine the relative strength of structural diversity (Gini index) in determining carbon stocks. Previous studies have highlighted that forest structure as tree density (e.g. Lewis *et al*., 2013; Porter *et al*., 2015; Zekeng *et al*., 2020) and basal area (van der Sande et al., 2017) drive biomass, and hence in this study we decided to test the effect of structural diversity in carbon stocks. Therefore, for each tree size group and the whole tree community per plot, the diversity Gini index was calculated using the Gini coefficient of tree basal area (Appendix S1). The Gini measures the inequality among values of tree size distribution. A Gini of zero expresses perfect equality while Gini of 1 expresses maximal inequality (i.e. high structural diversity) among values of the tree size distribution (Weiner & Solbrig, 1984).

### 2.6. Soil properties, topography and logging disturbance

The plots are situated in the region were topography is slightly uneven with a succession of low hills with generally gentle slopes (0.00-14.91%) interspersed with small well-marked streams and fine-scale variation in soil conditions. Therefore, per 1-ha plot five sampling points were used: one towards the four corners and one toward the center of the plot. More details about the collection and analyses of soil variables can be found in Appendix S2. Per sampling point, soil samples were taken between 0 and 20 cm for bulk density, texture, moisture content (MC), cations exchange capacity (CEC) and concentrations of carbon, total nitrogen (N_soil_), available phosphorus (P_soil_), the ratios between carbon and nitrogen (C:N_soil_) and nitrogen and phosphorus (N:P_soil,_ Appendix S3).

This study used four topographic variables: elevation, slope, curvature and aspect. Elevation was recorded throughout each 1-ha plot, at the four corners and the center and used to calculate topographic variables at the 1-ha scale. Mean elevation was calculated as the mean of the elevation measurements at the four corners and the center of a one ha plot. The slope was calculated as the average angular deviation from horizontal of each of the four triangular planes formed by connecting three of its four corners. Aspect is the direction of the slope faces, and cos (aspect) and sin (aspect) were calculated to make aspect data usable in linear models (Baldeck *et al*., 2013; Wang *et al*., 2017). Elevation and slope variables were obtained in the field during forest inventory using respectively GPS and clisimeter while the two other topographical variables were derived using ArcMap 10.1.

During field inventory, we found that some plots had been disturbed by logging which occurred 20 years ago, and hence we could not directly account the tree damage due to logging. Therefore, to take into account the entire disturbance (i.e. logging + damage), we measured the stumps of trees logged which were still present in the field and quantified it basal area using the empirical relationship equation: 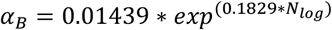 (Durrieu de Madron et al., 1998 but see Picard et al., 2012), where α_*B*_ is the proportion of damaged basal area and N_log_ the number of trees logged. Logging disturbance was computed as a continuous disturbance variable because logging disturbance depends on the distribution and density of commercial species and is therefore not evenly distributed in space and not varies strongly within the plot and between plots (Appendix S4). The relative logging disturbance (in %) was computed per ha, based on the basal area of all the stumps trees that were logged + damaged basal area divided by the total pre-logging basal area of the plot.

### 2.7. Statistical analyses

Structural Equation Modeling (SEM) offers the possibility to test multivariate and hierarchical direct and indirect relationships among the measured variables (Shipley, 2016). Therefore, we examined both direct effects, and indirect abiotic effects (topography, soil conditions and logging disturbance) on AGC stocks through taxonomic diversity and diversity Gini. We did this separately for small stems, understorey trees, large trees and the whole tree community (Fig. 1). Because we could also have many interactions among the predictive variables (e.g. the topographic variables can influence soil texture or soil fertility), we limited the number of possible models and the number of explanatory variables per model by evaluating only the framework corresponding to our *a priori*, as simple as possible hypothesis (see Fig. 1).

Because we had multiple indicator variables for abiotic conditions (i.e. soil and topographic variables; Appendix S3) and for taxonomic diversity, we first performed subsets regression analyses, including all topographic, soil fertility/texture, and taxonomic as the predictor variable and aboveground carbon stocks as a response variable. However, we included also disturbance and diversity Gini index in subsetting analyses to have their relative importance. From these results, we selected one for topographic and taxonomic variables or two variables for soil (i.e. one textural and one fertility variable) with the highest relative importance value (see Appendix S5). Soil variables were represented by soil texture (i.e. the proportion of clay, silt and sand) and soil fertility (i.e. CEC, C:N_soil_, EC, MC, N:P_soil_, N_soil,_ P_soil,_ pH), topography was represented by five variables (sine and cosine of aspect, elevation, terrain curvature, and terrain slope), and taxonomic diversity was represented by three variables (richness, rarefied species richness, and Shannon-Weaver index). Disturbance and structural diversity were included in all SEMs. Then per carbon stock variable, several SEMs were tested, from which we selected the SEM with the highest explained variation (*R*^*2*^) of the carbon stocks. The overall fit of the SEMs was assessed using χ^2^ –test (a *p*-value > 0.05 would indicate an absence of significant deviations between data and model, and means that the model is not rejected). In addition to the SEMs, simple bivariate relationships between biotic factors, abiotic factors and the carbon stocks variables using Spearman correlations showed that there is no collinearity between each group of factors (Appendix S7).

All analyses were performed in R 3.5.1. Correlations were evaluated using the *rcorr* function of the *Hmisc* package, linear mixed models with the lme function of the *nlme* package, and structural equation models with the *sem* function of the *lavaan* package (Rosseel, 2012).

## 3. Results

To evaluate the direct and indirect independent effects of abiotic and biotic variables on carbon stocks of different tree size groups and the whole tree community (Fig. 1), we used structural equation modeling (SEM). Only one model per carbon stocks variable was selected that was accepted by the Chi-square test and had the highest R^2^ for aboveground carbon stock (Fig. 2; Table 1) (see Appendix S8). The variation explained in carbon stocks ranged from 43% for the whole tree community to 72% for understorey trees (Fig. 2).

**Table 1.** The direct and indirect standardized effects of abiotic and biotic factors on carbon stock of all trees size classes (i.e. small stems, understorey trees and large trees) and the whole trees community level based on structural equation models (SEM; Fig. 2). The path coefficients (Coeff), standardized path coefficients (Std. coeff), Z-values and p-values are given for all regressions (i.e. all arrows in Fig. 2). All four models were accepted (p =0.06, 0.14, 0.59, 0.10 and χ^2^=3.65, 5.40, 0.29, 2.64 for carbon stocks of small stems, understorey trees, large trees and the whole tree community respectively; Appendix S8). Significant effects are indicated in bold (p < .05).

| SEM response variable | SEM predictor variable | Coeff | Std.Coeff | z-value | p-value |
| --- | --- | --- | --- | --- | --- |
| **Small stems** |  |  |  |  |  |
| Aboveground carbon stock (AGC) | Slope | 0.03 | 0.25 | 1.69 | 0.092 |
|  | **Nsoil** | **1.07** | **0.31** | **2.02** | **0.044** |
|  | Disturbance | 0.03 | 0.18 | 1.20 | 0.232 |
|  | Shannon-Weaver index | -0.04 | -0.03 | -0.22 | 0.825 |
|  | **Diversity Gini index** | **2.56** | **0.56** | **3.80** | **0.000** |
| Shannon-Weaver index | Slope | -0.02 | -0.15 | -0.80 | 0.425 |
|  | Nsoil | 0.35 | 0.12 | 0.64 | 0.520 |
|  | Disturbance | -0.03 | -0.24 | -1.38 | 0.167 |
| Diversity Gini index | Slope | 0.00 | 0.09 | 0.49 | 0.621 |
|  | Nsoil | -0.16 | -0.20 | -1.12 | 0.263 |
|  | Disturbance | 0.01 | 0.30 | 1.81 | 0.070 |
| $R^2$ AGC | | 0.54 | | | |
| $R^2$ Shannon-Weaver index | | 0.88 | | | |
| $R^2$ Diversity Gini index | | 0.83 | | | |
| **Understorey trees** |  |  |  |  |  |
| Aboveground carbon stock | Elevation | 0.01 | 0.21 | 1.09 | 0.277 |
|  | **Clay proportion** | **0.07** | **0.38** | **2.30** | **0.022** |
|  | Disturbance | -0.01 | -0.04 | -0.23 | 0.821 |
|  | Rarefied species richness | -0.06 | -0.31 | -1.96 | 0.050 |
|  | Diversity Gini index | 4.95 | 0.08 | 0.42 | 0.672 |
| Rarefied species richness | Elevation | -0.02 | -0.12 | -0.67 | 0.506 |
|  | Clay proportion | 0.17 | 0.19 | 1.05 | 0.295 |
|  | Disturbance | 0.09 | 0.05 | 0.27 | 0.784 |
| Diversity Gini index | Elevation | 0.00 | -0.31 | -1.83 | 0.068 |
|  | Clay proportion | 0.00 | 0.13 | 0.77 | 0.440 |
|  | Disturbance | 0.01 | 0.31 | 1.87 | 0.061 |
| $R^2$ AGC | | 0.72 | | | |
| $R^2$ Rarefied species richness | | 0.96 | | | |
| $R^2$ Diversity Gini index | | 0.8 | | | |
| **Large trees** |  |  |  |  |  |
|  | **Slope** | **5.22** | **0.40** | **3.20** | **0.001** |
|  | Clay proportion | 0.52 | 0.07 | 0.53 | 0.596 |
| Aboveground carbon stock | Disturbance | -3.66 | -0.26 | -1.91 | 0.056 |
|  | **Rarefied species richness** | **1.12** | **0.29** | **2.24** | **0.025** |
|  | **Diversity Gini index** | **868.59** | **0.58** | **4.27** | **0.000** |
| Rarefied species richness | Slope | -0.82 | -0.25 | -1.42 | 0.155 |
|  | Clay proportion | 0.34 | 0.18 | 1.03 | 0.305 |
|  | Disturbance | 0.77 | 0.21 | 1.21 | 0.227 |
| Diversity Gini index | Slope | 0.00 | 0.03 | 0.19 | 0.846 |
|  | **Clay proportion** | **0.00** | **0.35** | **2.12** | **0.034** |
|  | **Disturbance** | **0.00** | **0.35** | **2.11** | **0.035** |
| R<sup>2</sup> AGC |  | 0.43 |  |  |  |
| R<sup>2</sup> Rarefied species richness |  | 0.88 |  |  |  |
| R<sup>2</sup> Diversity Gini index |  | 0.8 |  |  |  |
| **Whole tree community** |  |  |  |  |  |
| Aboveground carbon stock | **Slope** | **4.67** | **0.36** | **3.02** | **0.003** |
|  | Clay proportion | 0.06 | 0.01 | 0.06 | 0.951 |
|  | Disturbance | -3.12 | -0.22 | -1.69 | 0.091 |
|  | **Species richness** | **0.91** | **0.33** | **2.60** | **0.009** |
|  | **Diversity Gini index** | **838.00** | **0.56** | **4.23** | **0.000** |
| Species richness | Slope | -0.36 | -0.08 | -0.44 | 0.657 |
|  | **Clay proportion** | **1.05** | **0.39** | **2.29** | **0.022** |
|  | Disturbance | 0.45 | 0.09 | 0.51 | 0.611 |
| Diversity Gini index | Slope | 0.00 | 0.03 | 0.19 | 0.846 |
|  | **Clay proportion** | **0.00** | **0.35** | **2.12** | **0.034** |
|  | **Disturbance** | **0.00** | **0.35** | **2.11** | **0.035** |
| R<sup>2</sup> AGC |  | 0.43 |  |  |  |
| R<sup>2</sup> Species richness |  | 0.85 |  |  |  |
| R<sup>2</sup> Diversity Gini index |  | 0.80 |  |  |  |

**Fig 2.**
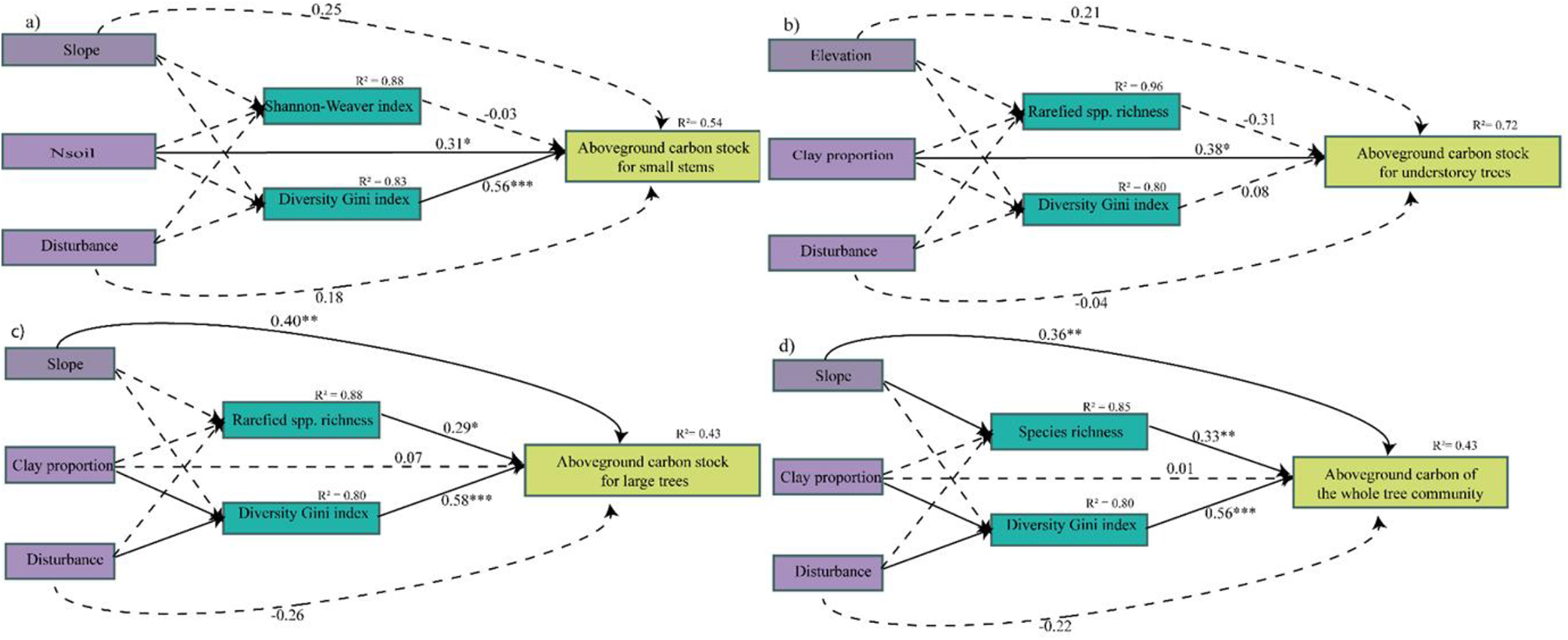
Structural equation models for the effects of the abiotic (i.e. topographic variables, soil fertility/texture and disturbance) and biotic variables (i.e. taxonomic richness and diversity Gini) on aboveground carbon stocks of the small stems (a), of the understorey trees (b), of the large trees (c) and of the whole tree community (d). All four models were accepted (Appendix S8). For all significant relationships (continuous black lines), the standardized regression coefficients and significance level are given only for direct relationships (*p <.05; **p <.01; ***p <.001), and for all non-significant relations (black, dashed lines), no statistics are shown. R^2^ values show the explained variance of the biotic factor and carbon stocks. More statistics of the structural equation model can be found in Table 1.

Different abiotic and biotic variables were selected per tree size group and the whole tree community in the SEMs. As a taxonomic diversity variable, the Shannon-Weaver index was selected for aboveground carbon (AGC) of small stems, rarefied species richness for AGC of understorey and large trees, and species richness for AGC of the whole tree community. Elevation was selected as a topographic variable influencing carbon stocks of understorey trees, and the terrain slope was selected for the two other trees size groups and the whole tree community. For soil variables, clay proportion was selected for AGC of the whole tree community and all tree size groups, except for small stems where Nsoil was selected (see Appendix S5 for results of all subsets regression analyses).

Biotic factors had generally strong and significant effects on AGC stocks, with 5 (63%) from the eight tested relationships being significant (Fig. 2; Table 1). The effects of taxonomic diversity were significant and positive on AGC of both large trees (*β* = 0.29; *p* = 0.03) and of the whole tree community (*β* = 0.33; *p* = 0.009), while effects of structural diversity (Gini index) was significantly positive for all AGC stocks except AGC of understorey trees (Fig. 2a, c and d; Table 1).

Abiotic factors had direct and indirect effects on AGC stocks (Fig. 2; Table 1). The terrain slope had a direct and significate positive effect only on AGC of large trees carbon (*β* = 0.40; *p* = 0.001) and the whole tree community (*β* = 0.36; *p* = 0.003). Elevation had a direct and non-significative positive effect on AGC of understorey trees (*β* = 0.21). Moreover, indirectly through taxonomic diversity and diversity Gini index, we found no significant effects of terrain slope and elevation on AGC of the whole tree community as well as all tree size groups (Table 1).

For soil variables, we found that Nsoil had a significant direct and positive effect on AGC for small stems (*β* = 0.31; *p* = 0.04). Soil texture (Clay proportion) had a direct and significant positive effect on AGC of understorey trees (*β* = 0.38; *p* = 0.02). Except, on AGC for small stems, via taxonomic diversity, we found that clay proportion increased AGC stocks. The same patterns were found also for the diversity Gini index effect (Fig. 2b, c and d; Table 1). The results showed that effects of clay proportion were significant and positive via species richness only on AGC for the whole tree community (Fig. 2d; Table 1) and via diversity Gini index on AGC of large trees and the whole tree community (Fig. 2c and d; Table 1).

We did not find any direct significant effect of logging disturbance on AGC of the whole tree community and the tree size groups (Fig. 2; Table 1). However, it indirectly increased AGC of large trees and the whole tree community through the diversity Gini index (Fig. 2c and d; Table 1).

## 4. Discussion

We asked how abiotic (topographic variables, soil variables and disturbance) and biotic variables (taxonomic and structural diversity) drive AGC stocks across trees size groups and the whole tree community and used structural equation modeling to test for their independent effects. We found that slope increased AGC of the whole tree community and the large trees while Nsoil and clay proportion increased AGC of the small stems and understorey trees groups. We also found that biotic factors have some indirect and direct effects on AGC. Here, we discuss the underlying mechanisms and the implications for the conservation, sustainable forest management and climate change mitigation potential of tropical forests.

### 4.1. Species diversity increases carbon stocks

We expected that taxonomic diversity (i.e. rarefied species richness, Shannon-Weaver index and species richness) would have a positive effect on AGC stock through niche complementary, the selection effect, and/or facilitation. However, we found that taxonomic diversity strongly drives carbon stock for only large trees and the whole tree community. The benefits of plant-plant interactions such as facilitation may explain these results. Hence some species could enhance soil fertility for the productivity of other species. But it might also be well possible that increasing species richness increases the chances of inclusion of highly productive favored dominant species (Ruiz-Benito *et al*., 2014). To our knowledge, this is the first local scale study analyzing the relationship between carbon stocks across trees size groups and the whole tree community of Cameroon tropical rainforest and its multiple underlying drivers. Most empirical studies that have examined the effects of diversity on forest carbon or productivity have ignored the effect of forest structure and environment (but see Day *et al*., 2013). The significant positive effects of rarefied species richness and species richness on aboveground carbon stocks of the large trees and the whole tree community found in this study have also been reported in several local and global tropical forest ecosystems (Cavanaugh *et al*., 2014; Con *et al*., 2013; Day *et al*., 2013; Poorter *et al*., 2017; van der Sande *et al*., 2018).

### 4.2. Structural diversity increased aboveground carbon stocks

We found a significant positive effect of structural diversity (Gini index) on aboveground carbon stocks of the whole tree community, large trees and understory trees (Fig. 2). When structural diversity is high, there is strong layering within the canopy which can more efficiently fit high amounts of biomass in the same area. Also, high structural diversity may indicate the presence of some very large trees that contribute disproportionally to forest biomass and carbon. This is confirmed by an earlier study in this forest, which showed that aboveground carbon at the whole tree community and at the large trees group are strongly driven by big-diameter trees (Zekeng *et al*., 2020).

### 4.3. Topography, soil conditions and disturbance shape aboveground carbon stocks

We expected that topographic variables could strongly affect carbon stocks. However, we found that topography (i.e., terrain slope) only affected AGB of large trees and of the whole tree community, while fertile soils only increased AGC of small stems (through Nsoil) and of understory stems (through clay content). The positive effect of slope on AGC of large trees and the whole tree community, showed evidence that differences in AGC stocks can result from topological constraints, particularly difference in terrain slope (Chave *et al*., 2003; de Castilho *et al*., 2006; Mensah *et al*., 2016; Salinas-Melgoza *et al*., 2018). It is important to note that dominant terrain slopes in our rainforest vary from 3 to 15%, and hence considered as steep slopes (Zare Chahouki *et al*., 2012). Aboveground carbon is expected to decrease in steep slopes because they have shallow soils (Gong *et al*., 2008), and are richer in the sand but poorer in silt content (Pachepsky *et al*., 2001), and hence are more vulnerable to erosion but surprisingly, we found out that AGC was higher on steeper slopes in our plots.

Soils of the semi-deciduous communal forest of Doume are leached, and hence may be nutrient-poor habitats. We therefore expected that increasing soil resources would strongly determine carbon storage. Soil nitrogen indeed significantly increased aboveground carbon stocks of small stems (Fig. 2a), and higher soil clay content – which is generally correlated with higher fertility – increased AGC of understory trees and structural diversity. It has been recognized as we found that soil textural properties are the most important characteristics of the soil, influencing, directly and indirectly, cascades of relations between soil nutrients, ions and soil drainage (Silver *et al*., 2000), and hence expected to have strong effects on AGC stocks. These results are in line with other studies (Lewis *et al*., 2013; van der Sande *et al*., 2018; Zarin *et al*., 2001), and demonstrate the importance of small-scale variation in soil conditions for the forest’s capacity to store carbon.

We did not detect any significant direct effects of logging disturbance on carbon stock of all tree size groups and of the whole tree community, maybe because it depends on the distribution of commercial species and they didn’t strongly vary between plots (Appendix S5). Contrary to our expectation, logging as a continuous variable did not reduce carbon stocks of the whole tree community, and all tree size groups (Fig. 2). These results may be due to the small variation in disturbance intensity and the time elapsed since the disturbance that has allowed carbon stocks to recover. Therefore, carbon stocks can rapidly recover. Contrary, disturbance increase significantly carbon stocks of large trees and the whole tree community through the diversity Gini index. It has been shown in the Amazonian forest that disturbance resulted in a decrease in AGB, but with time, it increases the recruitment of small trees (Holm *et al*., 2014) and hence this phenomenon could explain the results observed in our study. Our results, in conjunction with recent studies across Neotropical forest (Poorter *et al*., 2017; van der Sande *et al*., 2018) indicates that disturbance is an important process, by increasing the availability of light and others resources, hence promote the recruitment of small trees in the lower forest strata.

## 5. Conclusions and implications for carbon and REDD+

What is interesting in this study is not that AGC stocks are related to biotic and abiotic factors (that was expected) but how they are related within a Cameroonian tropical rainforest. The results showed that structural diversity has significant effects on aboveground carbon stocks of the whole tree community, large trees, and understory trees, which means that it is important to maintain a layered structure and also tall trees in the forest. We also found that aboveground carbon increased with increasing species richness and hence conserving biodiversity is not just an objective in itself. This result showed implication for REDD+ that forests with high diversity also tend to have high carbon stocks, indicating that forests with high carbon storage potential also have high conservation potential. Species richness could also help protect ecosystem productivity from environmental change (Isbell *et al*., 2011) and enhance the resilience of these ecosystems to disturbance (Díaz *et al*., 2009). Therefore, as diversity co-determines the functioning of the forest, many authors recommended that biodiversity conservation should not be seen as a simple simultaneous benefit of REDD+, but as integral and crucial components of all its activities (Díaz *et al*., 2009). Hence, due to his essential role in the forest functioning, biodiversity conservation is a win-win strategy for programs such as REDD+ and those under the Convention of Biological Diversity.

## Author’s contribution

JCZ and MMM.A designed the research; JCZ and PAE collected the data; JCZ. analyzed the data; JCZ and MS wrote the paper, and all the authors discussed the results and provided comments.

## Acknowledgments

The lead author is grateful for the Ph.D. exchange scholarship given by the Transdisciplinary Training for Resource Efficiency and Climate Change Adaptation in Africa II (TRECCAF-RICA II) project which is funded by the European Union. The research leading to these results has received financial funding from the British Ecological Society (EA17/1005), the Rufford Foundation (grant agreement N° 24895-1), and field material funding from the IDEA WILD Foundation. Masha van der Sande is supported by the Rubicon research program with project number 0.19.171LW.023, which is financed by the Netherlands Organisation for Scientific Research (NWO). We are grateful to the Conservation and Sustainable Natural Ressources Management Network (CSNRM-Net) Association for their logistical and technical support during the entire study. The authors thank all the technicians especially Mr. Oreeditse Kgosidintsi and Macpherson T Kavouras of the soil laboratory in the Department of Environmental Science at the University of Botswana for all their technical support during soil analyses. We are also grateful to the Doume municipality for their logistical support during the fieldwork. Specifically, we thank the mayoress and the secretary of the Doume municipality Mrs Mpans Giselle Rose and Ayinda Yannick respectively for their administrative diligence and for providing us with field permits. We furthermore, express our thanks to all those involved in fieldwork and data collection as well as community members of the different village of Doume.

