## Supplemental information for "Environmental, structural and taxonomic diversity factors drive aboveground carbon stocks in a semi-deciduous tropical rainforest strata in Cameroon"

Appendix S1. Descriptive statistics for biotic variables and the response variable (aboveground carbon stocks across trees size groups, and the whole tree community as response variables). Statistic (Mean, standard error's: SE, Minimum: Min, Maximum: Max) values are given across the 1-ha plots.

| | Variables | Unit | Mean | SE | Min | Max |
|---|---|---|---|---|---|---|
| Small stems (n=3.75 ha) | *Biotic factors* | | | | | |
| | Rarefied species index | | 1.64 | 0.04 | 1.00 | 2.00 |
| | Shannon-Weaver index | bit | 2.79 | 0.05 | 2.23 | 3.25 |
| | Species richness | | 24.37 | 0.88 | 14.00 | 32.00 |
| | Diversity Gini index | | 0.36 | 0.01 | 0.23 | 0.55 |
| | *Response variable* | | | | | |
| | Aboveground carbon | Mg $ha^{-1}$ | 1.63 | 0.23 | | |
| Understorey trees (n=15 ha) | *Biotic factors* | | | | | |
| | Rarefied species index | | 33.19 | 0.71 | 24.62 | 44.30 |
| | Shannon-Weaver index | bit | 3.43 | 0.05 | 2.70 | 3.81 |
| | Species richness | | 51.63 | 1.66 | 30.00 | 69.00 |
| | Diversity Gini index | | 0.20 | 0.002 | 0.17 | 0.23 |
| | *Response variable* | | | | | |
| | Aboveground carbon | Mg $ha^{-1}$ | 2.80 | 0.14 | 1.21 | 4.97 |
| Large trees (n=30 ha) | *Biotic factors* | | | | | |
| | Rarefied species index | | 103.62 | 1.54 | 80.70 | 119.95 |
| | Shannon-Weaver index | bit | 3.99 | 0.19 | 3.38 | 4.36 |
| | Species richness | | 104.83 | 2.18 | 75.00 | 124.00 |
| | Diversity Gini index | | 0.65 | 0.004 | 0.60 | 0.70 |
| | *Response variable* | | | | | |
| | Aboveground carbon | Mg $ha^{-1}$ | 177.61 | 6.11 | 114.05 | 231.49 |
| Whole tree community (n=30 ha) | *Biotic factors* | | | | | |
| | Rarefied species index | | 106.38 | 1.62 | 81.17 | 122.74 |
| | Shannon-Weaver index | bit | 4.05 | 0.03 | 3.67 | 4.36 |
| | Species richness | | 121.03 | 02.19 | 95.00 | 140.00 |
| | Diversity Gini index | | 0.73 | 0.004 | 0.68 | 0.77 |
| | *Response variable* | | | | | |
| | Aboveground carbon | Mg $ha^{-1}$ | 180.55 | 6.15 | 115.63 | 234.89 |

### Appendix S2: Collection and calculation of CEC, nutrient concentrations and litter variables.

Soil samples were collected using a soil auger at five specifics locations of the one ha plot (Subplots S1 of the four corners and the center). For each point of soil collection soil cores were dug at 20 cm depth. All sample were bulked, crushed and all organic no-decomposed material (stubbles, roots, stems and rubbish) were removed. In addition, five soil cores were collected at five position to determine its bulk density. All the sample collected were stored in zip-lock bags under cool temperatures before they were shipped to the soil laboratory of the Department of Environmental Science at the University of Botswana. After determining the soil moisture, all the soil sample were brought to the air-dried condition and then ground to pass a 2 mm mesh sieve.

**Soil moisture content**

Soil moisture content (MC) were evaluated as follow:

Crucibles were cleaned and dried for 1 hour. We weigh the empty crucible well marked with the weight ($W_1$) was recorded; then the wet soil was put into the crucible and the weight ($W_2$) was recorded. The crucible containing moist sample was placed at an oven box at 100 ° C for 48 hours or more until a constant weight was achieved. The crucibles were cooled in a desiccator and the weight ($W_3$) of the room temperature was recorded. The crucibles were cooled in a temperature and temperature ($W_4$) of the room temperature was recorded.

The MC was calculated as a percentage of the dry soil weight as follow:

$$\%MC = \frac{W_2 - W_3}{W_3 - W_1} * 100 .$$

where: $W_1$= Weight of crucible (g); $W_2$ = Weight of moist soil + crucible (g); $W_3$ = Weight of dried soil + crucible (g).

**Cation Exchange Capacity**

The Cation Exchange Capacity (CEC) of the soil was done using the Ammonium Acetate at pH= 7 method. In this study we obtained the CEC by summing the values obtained from the individual determination of the four elements. $Ca^{2+}$. $Mg^{2+}$. $K^+$ and $Na^+$. We weighed 12.5 g of air-dried into a 100 ml plastics bottles well labeled. 25 ml of neutral of ammonium acetate solution was added and the bottles were tightly and put on the shaker. They were shook for one hour at a speed of about 150 rpm after which the sample was removed from the shaker and allowed to stand for one hour. The mixture were transferred to a Buchner funnel filtered with a moist Whatman filter paper no. 42 under gentle suction. We continued to leach the soil slowly with small quantities of ammonium acetate until approximately 100 ml of leachate was collected. The leachate was transferred into 100 ml volumetric flask. Blank solution constituted only of ammonium acetate solution was also done.

An Agilent 4100 Microwave Plasma Atomic Emission Spectrometer (Agilent Technologies. Melbourne. Australia) equipped with an Inert One Neb nebulizer and a double-pass glass cyclonic spray chamber (Agilent Technologies. Melbourne. Australia) was used in this study to determine the concentrations of four elements above mentioned.

**Available phosphorus**

The available Phosphorus in soil was analysed using the Ascorbic Acid color development method. A weighed amount (5 g) of the air-dried samples was put into a 100 ml plastic bottles and then. 50 ml of 0.5 M of sodium bicarbonate ($NaHCO_3$) was added. The mixture was shaken for 30 min and filter through Whatman no. 5. One ml of the soil aliquot was pipetted into a test tube and successively 8 ml of distilled water, 1 ml of color solution, and 0.5 ml of ascorbic acid solution were added into the test tube. At each time the test tube was shaken and after putting ascorbic acid we wait for 15 minutes.

In parallel, standard P stock solution 100 ppm was prepared by dissolving 0.439 g of $KH_2PO_4$ in distilled water and dilute to 100 ml volumetric flask. Then, the standard p working solutions was prepared by pipetting 0. 2.0. 4.0. 8.0 ml of 100 ppm standard p stock solution into four 100 ml volumetric flasks and the volume was make up with distilled water. The Standard P curve was prepared by pipetting 1 ml of 0. 2.0. 4.0 and 8.0 ppm standard p working solutions respectively in 4 test tubes and successively 8 ml of distilled water, 1 ml of color solution and 0.5 ml of ascorbic acid solution were added. At each time the test tube was shaken and after putting ascorbic acid we wait for 15 minutes. The color intensity after 15 minutes was measured using Shimadzu UV Spectrophotometer 1800 with 880 nm wavelength.

P(ppm) = Pppm found from the standart curve*Dilution factor;

$P(\%) = Pppm*10^{-4}$

Appendix S3. Descriptive statistics for all abiotic variables (soil fertility, disturbance and topography). Statistic (Mean, standard error's: SE, Minimum: Min, Maximum: Max) values are given across the 1-ha

| Variables | Unit | Indicator of | Mean | SE | Min | Max |
|---|---|---|---|---|---|---|
| Electric conductivity | µSiemens | Soil fertility | 134.64 | 6.12 | 89.30 | 224.00 |
| pH | | Soil fertility | 6.42 | 0.11 | 4.87 | 7.00 |
| Clay proportion | % | Soil texture | 8.38 | 0.81 | 1.46 | 18.36 |
| Sand content | % | Soil texture | 77.65 | 3.15 | 0.00 | 94.97 |
| Silt content | % | Soil texture | 13.97 | 3.00 | 3.57 | 97.28 |
| Moisture content | % | | 26.87 | 1.09 | 14.26 | 38.25 |
| Cation exchange capaty (CEC) | cmol/kg | Soil fertility | 6.73 | 0.06 | 6.26 | 7.49 |
| Phosphorus (Psoil) | % | Soil fertility | 0.01 | 0.001 | 0.00 | 0.03 |
| Nitrogen (Nsoil) | % | Soil fertility | 0.23 | 0.02 | 0.06 | 0.37 |
| N:Psoil ratio | | Soil fertility and nutrient limitations | 25.66 | 04.15 | 3.64 | 88.47 |
| C:Nsoil ratio | | Soil fertility and nutrient limitations | 25.93 | 3.54 | 7.51 | 84.09 |
| Disturbance | % | Light availability | 2.98 | 0.57 | 0.00 | 8.31 |
| Elevation | m | Topography | 702.43 | 4.77 | 640.00 | 754.76 |
| Slope | % | Topography | 3.08 | 0.46 | 0.00 | 14.91 |
| Curvature | ° | Topography | 2.08 | 4.42 | -60.50 | 56.50 |
| Sine aspect | | Topography | 0.13 | 0.12 | -0.99 | 1.00 |
| Cosine aspect | | Topography | -0.09 | 0.14 | -0.99 | 0.99 |

**Appendix S4:** Harvest intensity per plot: control (plot without logging), plots with reduced impact logging with 1 trees ha-1 (RIL1), with 2 trees ha-1 (RIL2) and with 3 trees ha-1 (RIL3). The minimum and maximum disturbance level for the 1-ha plots are given.

| | **Control** | **RIL1** | **RIL2** | **RIL3** |
|---|---|---|---|---|
| Number of trees removed per hectare | 0 | 1 | 2 | 3 |
| Min. disturbance (% BA removed) | 0 | 2.92 | 4.40 | 7.22 |
| Max. disturbance (% BA removed) | 0 | 5.69 | 8.22 | 8.31 |

Appendix S5. Results of all subsets regression analyses for aboveground biomass carbon for the whole trees community level, large trees, understorey trees and small stems aboveground biomass (i.e.. the response variable), followed by averaging of all possible models. Per model, multiple indices for soil fertility and trait composition were included. The one or two soil fertility indices and taxonomic indices with the highest relative importance value (i.e.. the variables in bold) were selected for further analyses using structural equation modeling. Furthermore, the standardized regression coefficient (Std. coeff), adjusted standard error (SEadj), z-value and P-value are given.

| **Variable response** | **Predictor variables** | **Coeff** | **Adjusted SE** | **z value** | **Pr(>\|z\|)** | **Imp.value** |
|---|---|---|---|---|---|---|
| Aboveground carbon stock of small stems | **Diversity Gini index** | **2.73** | **0.41** | **6.61** | **<0.001** | **1** |
| | **Cation exchange capacity** | **0.48** | **0.17** | **2.84** | **0.004** | **0.95** |
| | **Disturbance** | **0.04** | **0.02** | **1.85** | **0.065** | **0.62** |
| | **Soil total Nitrogen** | **1.41** | **0.81** | **1.75** | **0.080** | **0.62** |
| | Soil available Phosphorus | 13.29 | 7.63 | 1.74 | 0.082 | 0.52 |
| | Ration Nitrogen :Phosphorus | -0.01 | 0.00 | 1.60 | 0.109 | 0.51 |
| | **Shannon-Weaver index** | **-0.18** | **0.14** | **1.25** | **0.212** | **0.43** |
| | **Slope** | **0.03** | **0.03** | **1.25** | **0.211** | **0.40** |
| | Ratio carbone :Nitrogen | 0.00 | 0.00 | 1.34 | 0.182 | 0.36 |
| | Rarefied species richness | -0.01 | 0.01 | 0.70 | 0.486 | 0.32 |
| | Silt proportion | <0.00 | 0.01 | 0.84 | 0.402 | 0.30 |
| | Sand proportion | 0.00 | 0.01 | 0.34 | 0.735 | 0.28 |
| | Elevation | <0.00 | 0.00 | 0.73 | 0.464 | 0.27 |
| | Rarefied species richness | 0.12 | 0.14 | 0.88 | 0.379 | 0.27 |
| | Moisture content | -0.01 | 0.01 | 0.96 | 0.337 | 0.24 |
| | Curvature | -0.00 | 0.00 | 0.39 | 0.694 | 0.23 |
| | Sine aspect | 0.03 | 0.10 | 0.32 | 0.751 | 0.22 |
| | Cosine aspect | -0.02 | 0.08 | 0.22 | 0.830 | 0.21 |
| | Clay proportion | 0.01 | 0.01 | 0.56 | 0.573 | 0.20 |
| | pH | 0.06 | 0.12 | 0.45 | 0.655 | 0.19 |
| | Electric conductivity | 0.00 | 0.00 | 0.08 | 0.934 | 0.15 |
| Aboveground carbon stock of understorey trees | **Diversity Gini index** | **-0.11** | **3.19** | **3.64** | **<0.001** | **0.99** |
| | **Cation exchange capacity** | **0.95** | **0.55** | **1.74** | **0.082** | **0.57** |
| | **pH** | **0.49** | **0.33** | **1.50** | **0.134** | **0.46** |
| | **Elevation** | **0.01** | **0.01** | **1.36** | **0.173** | **0.44** |
| | Clay proportion | 0.05 | 0.04 | 1.34 | 0.182 | 0.42 |
| | **Disturbance** | **-0.08** | **0.07** | **1.18** | **0.240** | **0.34** |
| | Cosine aspect | 0.18 | 0.21 | 0.89 | 0.373 | 0.29 |
| | **Rarefied species richness** | **-0.02** | **0.03** | **0.80** | **0.425** | **0.28** |
| | Sand proportion | -0.01 | 0.03 | 0.56 | 0.577 | 0.28 |
| | Sine aspect | -0.18 | 0.24 | 0.74 | 0.458 | 0.26 |
| | Electric conductivity | -0.01 | 0.01 | 0.85 | 0.396 | 0.26 |
| | Nsoil | 1.88 | 2.19 | 0.86 | 0.391 | 0.26 |
| | Curvature | <-0.00 | 0.01 | 0.60 | 0.549 | 0.25 |
| | Shannon Weaver index | 0.02 | 0.37 | 0.64 | 0.524 | 0.25 |
| | Slope | 0.04 | 0.06 | 0.57 | 0.571 | 0.24 |
| | Psoil - | 18.15 | 27.75 | 0.65 | 0.513 | 0.23 |
| | Moisture content | -0.02 | 0.03 | 0.65 | 0.513 | 0.22 |

| | | | | | | |
|---|---|---|---|---|---|---|
| | Silt proportion | -0.01 | 0.03 | 0.26 | 0.797 | 0.21 |
| | Ratio C:N | 0.01 | 0.01 | 0.61 | 0.544 | 0.21 |
| | Ratio N:P | -0.01 | 0.01 | 0.48 | 0.632 | 0.21 |
| | Species richness | 0.00 | 0.01 | 0.27 | 0.790 | 0.20 |
| Aboveground carbon stock of large trees | **Diversity Gini index** | **910.47** | **207.64** | **4.39** | **<0.001** | **1** |
| | **Slope** | **4.65** | **2.54** | **1.83** | **0.067** | **0.64** |
| | **Clay proportion** | **2.76** | **1.54** | **1.80** | **0.072** | **0.60** |
| | **Nsoil** | **-115.65** | **77.82** | **1.49** | **0.137** | **0.47** |
| | **Rarefied species richness** | **0.64** | **1.20** | **0.53** | **0.598** | **0.42** |
| | Species richness | 0.89 | 1.02 | 0.88 | 0.379 | 0.32 |
| | Shannon-Weaver index | -57.85 | 45.35 | 1.28 | 0.202 | 0.30 |
| | Sand proportion | -0.77 | 1.48 | 0.52 | 0.604 | 0.29 |
| | Curvature | 0.15 | 0.28 | 0.53 | 0.596 | 0.24 |
| | Ratio N:P | -0.24 | 0.44 | 0.55 | 0.579 | 0.24 |
| | Cosine Aspect | 3.10 | 8.56 | 0.36 | 0.717 | 0.22 |
| | Electric conductivity | -0.13 | 0.21 | 0.63 | 0.529 | 0.22 |
| | Psoil | -691.24 | 1082.15 | 0.64 | 0.523 | 0.22 |
| | Elevation | -0.06 | 0.26 | 0.23 | 0.816 | 0.21 |
| | Ratio C:N | 0.22 | 0.38 | 0.58 | 0.561 | 0.21 |
| | Silt proportion | -1.09 | 1.71 | 0.61 | 0.539 | 0.21 |
| | Sine Aspect | -1.04 | 10.11 | 0.10 | 0.918 | 0.20 |
| | pH | -0.77 | 14.99 | 0.05 | 0.959 | 0.20 |
| | Cation Exchange Capacity | 3.97 | 23.30 | 0.17 | 0.865 | 0.18 |
| | Disturbance | -0.30 | 3.01 | 0.10 | 0.920 | 0.18 |
| | Moisture content | -0.36 | 1.18 | 0.30 | 0.761 | 0.18 |
| Aboveground carbon stock of the whole trees community level | **Diversity Gini index** | **0.08** | **0.03** | **3.18** | **0.001** | **0.98** |
| | **Species richness** | **1.08** | **0.60** | **1.78** | **0.075** | **0.69** |
| | **Slope** | **4.68** | **2.55** | **1.84** | **0.067** | **0.64** |
| | **Clay proportion** | **2.81** | **1.54** | **1.82** | **0.068** | **0.61** |
| | **Nsoil** | **-114.44** | **78.18** | **1.46** | **0.143** | **0.46** |
| | Sand proportion | -0.79 | 1.51 | 0.52 | 0.600 | 0.29 |
| | Rarefied species richness | -0.12 | 1.17 | 0.15 | 0.881 | 0.26 |
| | Shannon-Weaver index | -0.23 | 0.63 | 0.37 | 0.709 | 0.25 |
| | Ratio N:P | -0.26 | 0.44 | 0.59 | 0.558 | 0.24 |
| | Curvature | 0.15 | 0.29 | 0.51 | 0.613 | 0.23 |
| | Cosine Aspect | 3.29 | 8.61 | 0.38 | 0.702 | 0.22 |
| | Electricity conductivity | -0.13 | 0.21 | 0.63 | 0.526 | 0.22 |
| | Psoil | -701.90 | 1093.19 | 0.64 | 0.521 | 0.22 |
| | Elevation | -0.05 | 0.26 | 0.20 | 0.841 | 0.21 |
| | Ratio C:N | 0.23 | 0.38 | 0.61 | 0.544 | 0.21 |
| | Silt proportion | -1.08 | 1.73 | 0.63 | 0.532 | 0.21 |
| | Sine Aspect | -1.21 | 10.17 | 0.12 | 0.906 | 0.20 |
| | pH | -0.18 | 15.08 | 0.01 | 0.990 | 0.20 |
| | Disturbance | -0.32 | 3.03 | 0.11 | 0.916 | 0.18 |
| | Cation Exchange Capacity | 5.01 | 23.43 | 0.21 | 0.831 | 0.18 |
| | Moisture content | -0.38 | 1.18 | 0.32 | 0.747 | 0.18 |

Appendix S6

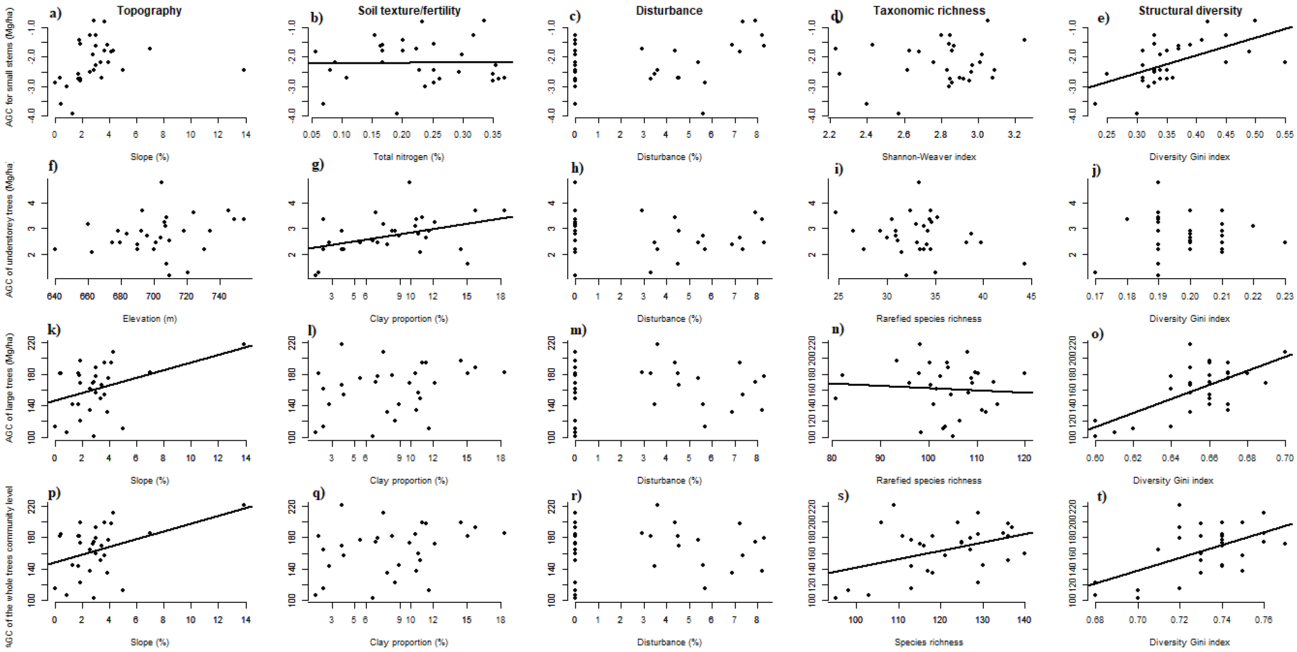


Figure S6. Bivariate relations of topography (a,f,k,p), soil texture/fertility (b,g,l,q), disturbance (c,h,m,r), taxonomic richness (d,i,n,s) and structural diversity (e, j,o,t) per 1-ha plot with aboveground carbon of small stems (a-e), aboveground carbon of understorey trees (f-j), aboveground carbon of large trees (k-o) and aboveground of the whole tree community (p-t). Each dot is a 1-ha plot.

Appendix S7:

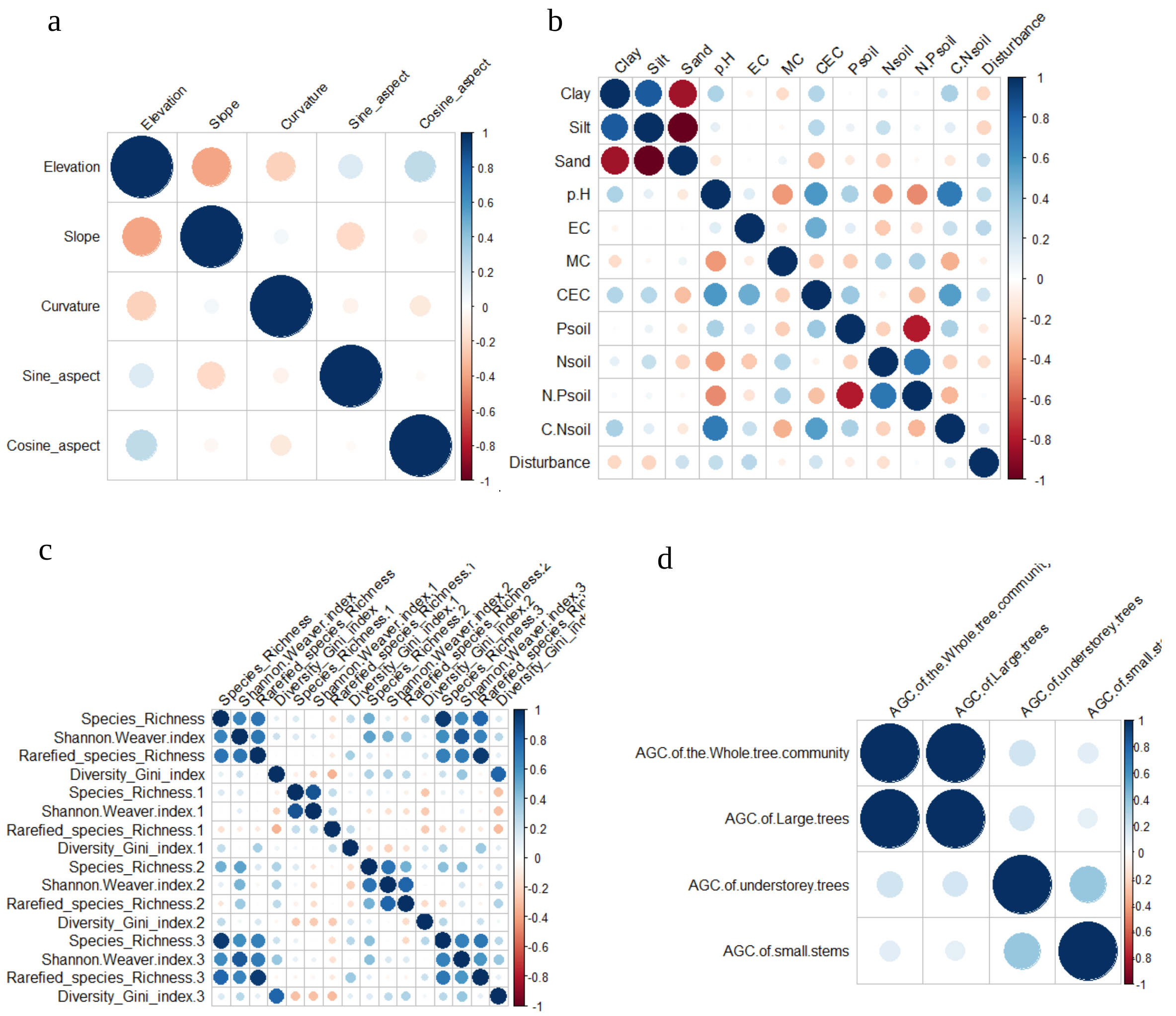


**Figure S7:** Spearman correlations between (a) topographic variables, (b) disturbance and soil variables, (c) Biotic variables (1 correspond to small stems, 2 equal to understorey trees, 3 equal to large trees and the variables without number represent variables for the whole tree community), and (d) aboveground carbon (AGC) for the trees size groups and the whole community. Dark blue colours indicate strong positive correlation and dark red colours indicate strong negative correlations. Topographic variables were non-significant and weak. Aboveground carbon of the whole tree community was significantly positively correlated only with AGC of large trees. Fine root biomass was significantly positively correlated with aboveground biomass and significantly negatively with aboveground biomass productivity, but the other correlations between stocks of biomass, fine roots, soil organic matter, and productivity were non-significant and weak. Soil variables were not significantly correlated, except for a logical strong negative correlation between $P_{soil}$ and $N:P_{soil}$ and also between sand and clay soil texture. For biotic variables, there is significant positive correlation only between species richness and rarefied species richness of the whole tree community and the same variables for large trees.

Appendix S8. Statistics showing the model fit of structural equation models (SEMs) for carbon stock for the whole tree community, large trees, understorey trees and small stems. A *P*-value > 0.05 indicates that the model is accepted.

| Response variables | Topographic variables | Taxonomic diversity variables | Soil variables | Model Chi-squared | Model *P*-value | $R^2$ of response variables |
|---|---|---|---|---|---|---|
| Aboveground carbon stock for the whole tree community | Slope | Species richness | Clay proportion | 2.642 | 0.104 | 0.43 |
| | | | $N_{Soil}$ | 0.854 | 0.356 | 0.42 |
| AGC of large trees | Slope | Rarefied species richness | Nsoil | 0.791 | 0.374 | 0.42 |
| | | | Clay proportion | 0.292 | 0.589 | 0.43 |
| AGC of understorey stems | Elevation | Rarefied species richness | Clay | 5.401 | 0.144 | 0.72 |
| | | | CEC | 4.372 | 0.224 | 0.64 |
| AGC for small stems | Slope | Shannon-Weaver index | Nsoil | 3.653 | 0.056 | 0.54 |
| | | | CEC | 2.763 | 0.096 | 0.33 |
